## Supplementary material for "Model of inverse bleb growth explains giant vacuole dynamics during cell mechanoadaptation": SI Appendix

**This PDF file includes:**

Supplementary text
Figs. S1 to S2
References for SI reference citations

### Supplementary Text

This *Supplementary Text* is organized as follows. In Section I, we characterize the mechanical equilibrium of the system after perfusion. We show that Laplace law holds for both the cell and the inverse bleb separately. The resulting equations, however, are not independent but are related through the intracellular pressure. We provide evidence that correction terms due to membrane bending are negligible for the giant vacuoles (GVs) typically observed *in vivo* in Schlemm's canal endothelial cells. Section II presents the linear stability analysis of the stationary configurations of giant vacuoles described by Eq. (3) (main text). In Section III we study numerically the growth and collapse of GVs after reducing the cortical tension $\sigma_c$ (e.g., by treating the cell with the Rho-kinase Y-27632).

### I. Mechanical equilibrium for the perfused system

In steady-state conditions, the mechanical equilibrium of the perfused system is specified as follows: (a) For the inflated cell, Laplace law holds. This relates the intracellular pressure to the cell radius as

$$P - P_e = \frac{2\sigma}{R} \tag{1}$$

with $\sigma$ the surface tension of the cell (Fig. S1 B). (b) For the inverse bleb, the corresponding mechanical equilibrium condition is determined by minimizing its free energy $F$ with respect to the independent variables $r$ and $\theta$. In complete generality, this free energy comprises (i) the surface energy of the inverse bleb $F_s$ (proportional to its area); (ii) the energy due to the perfusion pressure drop $F_p$ (proportional to its volume); (iii) the bending energy $F_b$ as specified by the Helfrich Hamiltonian (Helfrich, 1973); and (iv) a Lagrange multiplier $L$ imposing the geometrical condition $r\sin\theta = a$. Putting those terms into equations, we write

$$\begin{aligned} F &= F_b + F_s + F_p + L(r\sin\theta - a) \\ &= 2\kappa_m \int H^2\, dA + \Sigma \int d\,A - (p-P)\int dV + L(r\sin\theta - a) \end{aligned} \tag{2}$$

with $\kappa_{\mathrm{m}}$ the membrane bending modulus, $H = -1/r$ the mean curvature of the inverse bleb, $\Sigma$ its surface tension and $p - P$ the pressure drop at the cell-inverse bleb interface. Substituting area and volume of the inverse bleb, respectively $S = 2\pi r^2(1-\cos\theta)$ and $V = \frac{4\pi}{3} r^3 (2+\cos\theta)\left[\sin\left(\frac{\theta}{2}\right)\right]^4$, we obtain

$$\begin{aligned} F = 2\pi(1-\cos\theta)&\left\{2\kappa_{\mathrm{m}} + r^2\left[\Sigma - \frac{p-P}{3} r\big(1-\cos\theta\,(1+\cos\theta)\big)\right]\right\} \\ &+ L(r\sin\theta - a). \end{aligned} \tag{3}$$

The equilibrium configuration is obtained by solving $\partial_r F = 0 = \partial_\theta F$. In details,

$$\frac{\partial F}{\partial r} = 2\pi r(1-\cos\theta)\left[2\Sigma - (p-P)r\left(1 + \frac{1}{2}\cos\theta\,(1+\cos\theta)\right)\right] + L\ \sin\theta, \tag{4}$$

$$\frac{\partial F}{\partial \theta} = 4\pi\kappa_{\mathrm{m}} \sin\theta + 2\pi r^2 \sin\theta \left[\Sigma - \frac{(p-P)r}{2}(\sin\theta)^2\right] + Lr\cos\theta. \tag{5}$$

Setting these equations to zero yields the equilibrium condition

$$p - P = \frac{2\Sigma}{r} + \frac{4\kappa_{\mathrm{m}}}{r^3}\frac{1+\cos\theta}{1-\cos\theta}. \tag{6}$$

The free energy of the inverse bleb is thus obtained by substituting Eq. (6) into Eq. (3). We then obtain

$$F = \frac{2\pi}{3}\Sigma r^2 - \frac{2\pi}{3}\Sigma r^2\left(1 - \frac{a^2}{r^2}\right)\cos\theta + 4\pi\kappa_{\mathrm{m}}\left(1 - \frac{2}{3}\frac{a^2}{r^2}\right) - 4\pi\kappa_{\mathrm{m}}\left(1 + \frac{a^2}{3r^2}\right)\cos\theta. \tag{7}$$

Inverse bleb configurations exist for all $r$ because $F \geq 0$ (recall that $\theta \in [\pi/2, \pi)$).

We now show that the bending term in the rhs of Eq. (6) can be effectively neglected. The contribution due to bending is expected to be negligible for sufficiently large $r$, such that we can use the expansion $\cos\theta \simeq 1 - a^2/(2r^2)$. Substituting this expression into Eq. (6), we can write

$$p - P = \frac{2\Sigma}{r} + \frac{4\kappa_{\mathrm{m}}}{r^3}\frac{1-|\cos\theta|}{1+|\cos\theta|} \simeq \frac{2\Sigma}{r}\left(1 + \frac{\kappa_{\mathrm{m}}}{2\Sigma}\frac{a^2}{r^4}\right), \tag{8}$$

where the second term in parentheses satisfies $\kappa_m a^2/(2\Sigma r^4) \lesssim \kappa_m/(2\sigma_0 r^2)$. This term can thus be neglected when $\kappa_m/(2\sigma_0 r^2) \lesssim 1$, which yields $r \gtrsim \sqrt{\kappa_m/(2\sigma_0)}$. For typical values of $\kappa_m$ and $\sigma_0$ ($\sigma_0 \simeq 414$ pN/$\mu$m (Tinevez, et al., 2009); $\kappa_{\mathrm{M}} \simeq 43$ pN $\cdot$ nm (Phillips, Kondev, Theriot, & Garcia , 2012)), this yields $r \gtrsim 0.007\mu$m, which is satisfied for the inverse blebs considered here (in fact $r \geq a$).

Therefore, the bending term in Eq. (6) can be neglected. The mechanical equilibrium of the inverse bleb is thus described by the Laplace law

$$p - P = \frac{2\Sigma}{r}. \tag{9}$$

### III. Linear stability analysis of the stationary configurations of giant vacuoles

The linear stability of the solutions of Eq. (3) (main text) is studied by computing the radial force

$$f = p - P - \frac{2\Sigma}{r}, \tag{10}$$

which is exerted on the surface of the GV in response to a small positive perturbation $\delta r$ of its radius. Positive force indicates an unstable configuration; vice versa, negative force indicates a stable configuration.

We call $(r^*, \theta^*, R^*, P^*, \bar{\sigma}^*)$ the stationary configuration of the perfused system obtained by solving Eq. (3) (main text) complemented with volume conservation and a suitable functional prescription for $\bar{\Sigma}$ and $\bar{\sigma}$ (main text, Fig. 3). We now consider a small positive perturbation $\delta r$ of the vacuole radius and denote the perturbed solutions accordingly as $r = r^* + \delta r$, $\theta = \theta^* + \delta\theta$, $R = R^* + \delta R$, $P - P_e = P^* + \delta P$, $\sigma = \bar{\sigma}^* + \delta\sigma$ and $\Sigma = \bar{\Sigma}^* + \delta\Sigma$. Vacuole and cell perturbed surface tension are given by

$$\Sigma = \bar{\Sigma} + 2E_C h_C\left(1 - \frac{r^*}{r}\right), \qquad \sigma = \bar{\sigma}, \tag{11}$$

where for the inverse bleb we also consider the elastic resistance to deformations due to additional contractile actomyosin structures (here assumed to envelop the bleb and of thickness $h_C$). The angular perturbation follows from the geometrical constraint, $r\sin\theta = a$, i.e.,

$$\theta = \pi - \sin^{-1}\left(\frac{a}{r^* + \delta r}\right) \simeq \pi - \sin^{-1}\left[\frac{a}{r^*}\left(1 - \frac{\delta r}{r^*}\right)\right] \simeq \theta^* + \frac{\sin\theta^*}{|\cos\theta^*|}\frac{\delta r}{r^*}. \tag{12}$$

Using this expansion, we find

$$\cos\theta \simeq \cos\left(\theta^* + \frac{\sin\theta^*}{|\cos\theta^*|}\frac{\delta r}{r^*}\right) \simeq \cos\theta^* - \frac{\left(\sin\theta^*\right)^2}{|\cos\theta^*|}\frac{\delta r}{r^*}. \tag{13}$$

Volume conservation yields the cell radius perturbation. Expanding the equation $R^3 = R_0^3 + 2r^3(2 + \cos\theta)\sin^4(\theta/2)$ to first order in $\delta r, \delta R$, we obtain

$$\delta R = \frac{r^{*2}}{4R^{*2}}\left(1 - \cos\theta^*\right)^2\left[2 + \cos\theta^* + 2\frac{\left(1 + \cos\theta^*\right)^2}{|\cos\theta^*|}\right]\delta r. \tag{14}$$

We note that this quantity is positive for all $\theta^*$. Similarly expanding Eqs. (11), we find the perturbation of the surface tensions, i.e.,

$$\delta\Sigma = \bar{\Sigma}'^*\delta S + \frac{2E_C h_C}{r^*}\delta r, \qquad \delta\sigma = \bar{\sigma}'^*\delta A, \tag{15}$$

with $\bar{\Sigma}'^* = \partial_S\bar{\Sigma}(S^*)$ and $\delta S$ the perturbation of its area; likewise, $\bar{\sigma}'^* = \partial_A\bar{\sigma}(A^*)$ and $\delta A$ the perturbation of its area. Area perturbations of inverse bleb and cell are given by

$$\delta S = 2\pi\left[2\left(1 - \cos\theta^*\right) + \frac{\left(\sin\theta^*\right)^2}{|\cos\theta^*|}\right]r^*\delta r, \qquad \delta A = 6\pi R^*\delta R, \tag{16}$$

both also positive for all $\theta^*$. For the intracellular pressure we expand the Laplace law $P - P_e = 2\sigma/R$, which yields

$$\delta P = \frac{2}{R^*}\delta\sigma - \frac{2\bar{\sigma}^*}{R^{*2}}\delta R. \tag{17}$$

The radial force exerted on the GV surface is thus given by

$$f \simeq -\delta P - \frac{2}{r^*}\delta\Sigma + \frac{2\bar{\sigma}^*}{r^{*2}}\delta r \tag{18}$$

where we employed the equilibrium relation $p - P^* - 2\bar{\Sigma}^*/r^* = 0$. Combining Eqs. (16), (17) and (18) finally yields

$$\frac{f}{\delta r} = \frac{2(\bar{\Sigma}^* - 2E_C h_C)}{r^{*2}} - \frac{2\bar{\Sigma}'^*}{r^*}\frac{\delta S}{\delta r} - \frac{2\bar{\sigma}'^*}{R^*}\frac{\delta A}{\delta r} - \frac{2\bar{\sigma}^*}{R^{*2}}\frac{\delta R}{\delta r} \tag{19}$$

We consider first the case when there are no additional contractile structures on the inverse blebs, i.e., $h_C = 0$. For configurations of inverse blebs in the cortex-dominated regime, the steady-state surface tensions of the inverse blebs and cell are assumed constant, $\bar{\Sigma}^* = \bar{\sigma}^* = \sigma_0$. As such, $\bar{\Sigma}'^* = \bar{\sigma}'^* = 0$. Eq. (19) thus reduces to

$$\frac{f}{\delta r} = \frac{2\sigma_0}{r^{*2}}\left(1 - \frac{r^{*2}}{R^{*2}}\frac{\delta R}{\delta r}\right) \tag{20}$$

This quantity is positive for all $r^*$; therefore, these inverse bleb configurations are unstable. By thinking in terms of dynamical systems theory, we expect configurations of inverse blebs in the membrane-dominated regime to be stable. However, there is no similar simple analytical argument to demonstrate this. We can then study numerically Eq. (19). Fig. S1A confirms that vacuoles in the cortex-dominated regime are unstable (force is positive); Fig. S1B indeed shows that vacuoles in the membrane-dominated regime are stable as expected (force is negative).

Eq. (19) also enables us to estimate the thickness of additional cortex that the cells needs to build to stabilize vacuoles in the cortex-dominated regime. Imposing the force to be negative, and solving with respect to $h_C$ we find

$$h_C \geq \frac{\sigma_0}{2E_C}\left[1 - \left(\frac{r^*}{R^*}\right)^2 \frac{\delta R}{\delta r}\right] \tag{21}$$

Recalling that $\delta R/\delta r \propto \left(\boldsymbol{r}^*/\boldsymbol{R}^*\right)^2$, the term in brackets is circa one. Therefore, we obtain the approximate expression $\boldsymbol{h_C} \gtrsim \boldsymbol{\sigma_0}/\boldsymbol{2}E_C \approx \boldsymbol{23}$ nm.

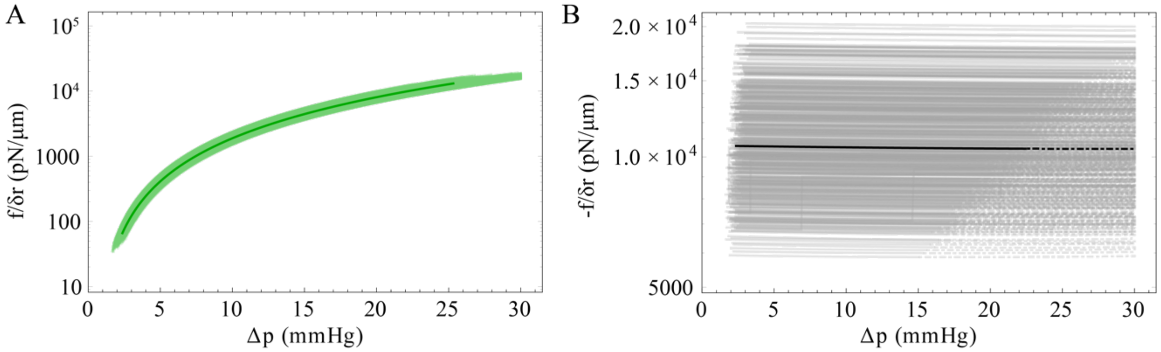


**Figure S1. Linear stability of giant vacuoles.** The linear stability of a giant vacuole against small positive perturbations of its radius is investigated by calculating numerically the response force exerted on the vacuole, Eq. (10). Reference curves (darker lines) correspond to the choice of model parameters (same as in Fig. 2B, main text): $R_0 = 10\ \mu$m, $a = 0.25\ \mu$m, $d = 10\ \mu$m, $\sigma_m = 40$ pN/$\mu$m, $\sigma_c = 374$ pN/$\mu$m, $\varepsilon^* = 0.5$ and $K_m = 10^5$ N/m. Lytic values of the surface tension (dot-dashed line) corresponds to a local relative areal strain 5% $\varepsilon^*$. Blurred regions correspond to values obtained by allowing for up to 20% variability in the parameters (according to Table 2, main text). A. Cortex-dominated regimes. B. Membrane-dominated regime.

#### III. Growth and collapse of giant vacuoles upon cortical tension impairment

In Fig. S2 we study numerically how the growth and collapse processes of GVs are affected by a reduction of the cortical tension $\sigma_c$. This can be realized experimentally, e.g., by treating Schlemm's canal endothelial cells with the Rho-kinase inhibitor Y-27632, which induces cell relaxation and disassembly of actin stress fibers and focal adhesions (thus reducing $\sigma_c$) (Vasantha, Deng, Kumar, & Epstein, 2001) (Honjo, et al., 2001) (Rosenthal, et al., 2005) by inhibiting the phosphorylation of the regulatory myosin light chain (Davies, Reddy , Caivano, & Cohen, 2000) (Ishizaki, et al., 2000) (Uehata, et al., 1997) (Kaibuchi, Kuroda, & Amano, 1999). Numerical simulations reveal: a slight decrease of the characteristic timescale for GV inflation (about $1.2-1.6$ min in this case versus about $1.6-2.2$ min for the wildtype); and a significant increase of the characteristic timescale for GV collapse upon removal of the pressure drop (about $4-6$ min for the wildtype versus about $8-10$ min for the treated case).

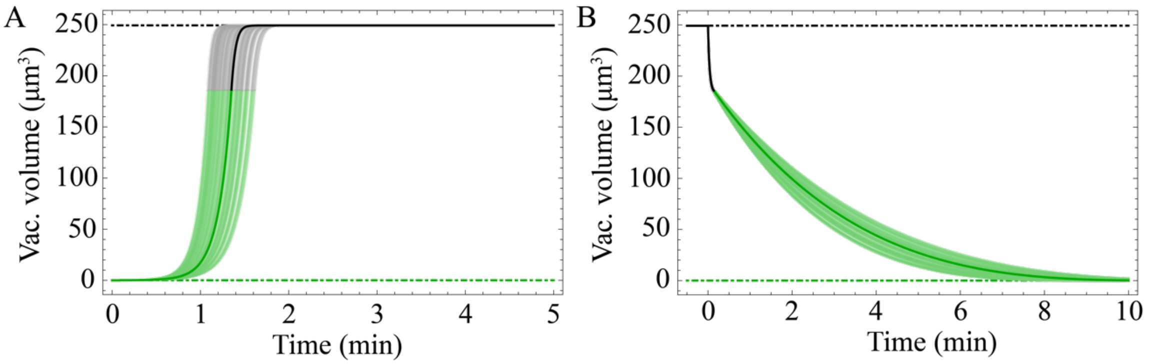


**Figure S2. Growth and collapse of giant vacuoles upon reduction of the cortical tension.** We set $\boldsymbol{\sigma}_c = \mathbf{187}$ pN/$\boldsymbol{\mu}$m. Pressure protocols, initial conditions and other model parameters are the same as in Fig. 4 **(**main text**)**. (A) Giant vacuole growth. The characteristic timescale for vacuole inflation is decreased compared to wildtype (about $\mathbf{1.6-2.2}$ min). (B) Giant vacuole collapse. The characteristic timescale for vacuole deflation is significantly increased compared to wildtype (about $\mathbf{4-6}$ min).
